# Behavioral metabolic suppression confounds thermal performance estimates and climate vulnerability assessments in a marine ectotherm

**DOI:** 10.64898/2026.07.13.738316

**Authors:** Cameron Edgar, Henry Penfold, Tanya Martinez, Christopher D. Wells

**Author notes:** Corresponding author: Christopher D. Wells.

## Abstract

Thermal performance curves (TPCs) predict species’ vulnerability to climate change, but standard respirometry assumes that measured oxygen consumption reflects physiological state. Sessile invertebrates that retract their tentacles and contract under thermal stress violate this assumption, with unmeasured consequences for thermal limit estimates.
We tested this behavioral confound in an undescribed cold-water intertidal anemone (*Urticina* sp.) in the Northwest Atlantic by integrating a negative binomial encounter-rate regression, a maximum entropy species distribution model (both from effort-corrected iNaturalist data), and closed-chamber respirometry across seven temperatures (1-30°C).
The strongest distributional predictors were cloud cover and coastal urbanization, with a weaker association with winter minimum SST; direct evidence for warm-edge thermal limitation came from the experiment. Anemone expansion state was variable and without a clear trend across the coldest treatments, declining above 20°C, and collapsing at the 30°C treatment, which proved lethal.
Standard TPC models extrapolated the thermal maximum far beyond the lethal bracket. A Bayesian multiplicative model that separated physiology from behavior showed that physiology continued to track temperature while expansion state declined above 20°C; a fully expanded anemone respired about twice as fast as a fully closed one at the same temperature. The decline in measured respiration is therefore both behavioral and physiological, and disentangling the two requires recording expansion state alongside oxygen consumption.
Because a closed anemone cannot feed or exchange gases, the ecologically relevant thermal limit is the temperature at which the animal can no longer maintain its normal expanded posture, not a curve-fitted thermal maximum. That behavioral threshold leaves warm-edge populations within a few degrees of functional thermal failure.
Future thermal physiology studies of organisms capable of modulating oxygen consumption through behavior should incorporate quantitative behavioral covariates to separate physiological from behavioral components of the metabolic response.

**Résumé:**

1. Les courbes de performance thermique (CPT) prédisent la vulnérabilité des espèces au changement climatique, mais la respirométrie classique postule que la consommation d’oxygène mesurée reflète l’état physiologique. Les invertébrés sessiles qui rétractent leurs tentacules et se contractent sous stress thermique ne satisfont pas cette hypothèse, avec des conséquences non quantifiées sur l’estimation des limites thermiques.
2. Nous avons testé ce facteur de confusion comportemental chez une anémone intertidale d’eau froide non décrite (*Urticina* sp.) de l’Atlantique Nord-Ouest, en combinant une régression binomiale négative du taux de rencontre, un modèle de distribution d’espèces à maximum d’entropie (tous deux sur données iNaturalist corrigées de l’effort) et une respirométrie en chambre close à sept températures (1 à 30 °C).
3. Les principaux prédicteurs étaient la couverture nuageuse et l’urbanisation côtière, l’association au minimum hivernal de température de surface de la mer étant plus faible. La preuve directe d’une limitation thermique en bordure chaude provient de l’expérience. L’état d’expansion variait sans tendance nette aux traitements les plus froids, diminuait au-delà de 20 °C et s’effondrait à 30 °C, traitement qui s’est révélé létal.
4. Les CPT classiques ont extrapolé le maximum thermique bien au-delà de l’intervalle létal. Un modèle bayésien multiplicatif séparant physiologie et comportement a montré que la physiologie suivait toujours la température alors que l’état d’expansion diminuait au-delà de 20 °C; à température égale, une anémone pleinement épanouie respirait environ deux fois plus qu’une anémone pleinement fermée. La baisse mesurée est donc à la fois comportementale et physiologique, et les dissocier exige d’enregistrer l’état d’expansion avec la consommation d’oxygène.
5. Une anémone fermée ne pouvant ni se nourrir ni assurer ses échanges gazeux, la limite thermique écologiquement pertinente est la température à laquelle l’animal ne peut plus maintenir son expansion normale, et non un maximum issu d’un ajustement de courbe. Ce seuil comportemental place les populations de bordure chaude à quelques degrés de la défaillance thermique fonctionnelle.
6. Les futures études de physiologie thermique sur des organismes capables de moduler leur consommation d’oxygène par leur comportement devraient intégrer des covariables comportementales quantitatives pour séparer les composantes physiologique et comportementale de la réponse métabolique.

## Introduction

Climate change is reshaping marine communities worldwide, with leading range edges of marine species expanding poleward at a mean rate of 72 km per decade (Poloczanska *et al*. 2013), several times faster than the 6-17 km per decade seen on land (Parmesan & Yohe 2003; Chen *et al*. 2011). Predicting which species will keep pace with warming and which will fail requires understanding how performance maps onto temperature. Thermal performance curves (TPCs) describe how fitness-related traits vary with temperature and define the limits at which performance collapses (Huey & Kingsolver 1989). From a TPC one can derive the thermal safety margin: the difference between an organism’s upper thermal limit and the warmest temperatures it currently experiences (Sunday *et al*. 2014). Species with narrow safety margins face the greatest risk of local extirpation due to warming (Deutsch *et al*. 2008; Pinsky *et al*. 2019). Sessile marine species that cannot relocate to cooler microhabitats are particularly vulnerable (Helmuth *et al*. 2002; Helmuth *et al*. 2006) with extirpations twice as common in the ocean as on land (Pinsky *et al*. 2019).

Whether those safety margins are accurate depends on whether the TPCs behind them actually capture physiological thermal sensitivity, and a growing body of evidence suggests they often do not (Halsey *et al*. 2015; Schulte 2015; Clusella-Trullas *et al*. 2021). Closed-chamber respirometry measures whole-organism oxygen consumption as a proxy for metabolic rate and assumes that changes in respiration track changes in physiological state (Killen *et al*. 2021). Three problems challenge that assumption. First, metabolic TPCs do not reliably represent whole-organism fitness: different performance traits within the same species peak at temperatures that can differ by nearly 7°C (Kellermann *et al*. 2019). Second, behavioral responses to temperature can be more thermally sensitive than the underlying physiology, so activity windows are often narrower than physiological tolerance ranges (Gunderson & Leal 2016). For example, in intertidal ectotherms, locomotion ceases well before physiological failure (Monaco, McQuaid & Marshall 2017). Third, behavior alone can produce dome-shaped TPCs even when there is no underlying shortfall between oxygen supply and demand (Neubauer & Andersen 2019), meaning that the declining half of a metabolic TPC may reflect an animal withdrawing rather than a cell failing. Sessile invertebrates that modulate their metabolism through posture exemplify the problem. Sea anemones contract under thermal stress, reducing the surface available for gas exchange and suppressing respiration independently of any change in cellular demand (Shick 1991).

Bivalves close their valves and depress metabolic rate to as little as 10% of their standard rate, with closure preceding molecular stress markers (Ortmann & Grieshaber 2003; Anestis *et al*. 2007). In each case, the organism suppresses its metabolic expression at ambient temperature, and standard respirometry measures the integrated product of this suppression and any physiological response. The relative contribution of each has never been partitioned from a single respirometry trial. No study has proposed a way to disentangle these contributions with a single fitted parameter scaling a measured behavioral covariate taken alongside oxygen consumption.

Rocky intertidal ecosystems provide an ideal system for quantifying this confound, because sessile invertebrates dominate the fauna and behavioral closure under thermal and desiccation stress is widespread. Intertidal organisms experience some of the most extreme thermal conditions in the marine realm, with body temperatures during low tide driven far above ambient sea surface temperatures (SSTs) by aerial exposure and solar radiation (Helmuth *et al*. 2002; Helmuth *et al*. 2006). Physical stress sets species’ upper distributional limits while biotic interactions constrain their lower boundaries (Connell 1972), making these communities indicators of climate-driven ecological change (Mieszkowska *et al*. 2006). The Northwest Atlantic highlights the urgency: the Gulf of Maine warmed faster than 99.9% of the global ocean between 2004 and 2013, with surface temperatures increasing at approximately 0.23-0.26°C per year since 2004 (Mills *et al*. 2013; Pershing *et al*. 2015). This rapid warming has restructured fisheries, altered deepwater circulation, and placed sessile invertebrates at particular risk because they cannot behaviorally track shifting isotherms (Harley 2011; Pershing *et al*. 2015; Record *et al*. 2019). Yet fundamental thermal physiology data remain absent for most intertidal invertebrates in this region.

Filling these gaps requires pairing distributional evidence with direct physiological measurement, and citizen science platforms now make the distributional component feasible. iNaturalist, with over 200 million observations as of September 2024, has become a leading data source for non-avian species distribution modeling (Mason *et al*. 2025), though its well-documented spatial and taxonomic biases require explicit correction (Di Cecco *et al*. 2021). Observer-oriented approaches that use co-occurring non-target taxa as proxies for sampling intensity (Milanesi, Mori & Menchetti 2020) allow bias-corrected distributional models to be built from opportunistic data. Paired with experimental TPCs, these distributional models can test whether the thermal gradient associated with a species’ geographic range also governs its physiological performance, and whether the methods used to characterize that performance are themselves reliable.

We propose that for sessile marine invertebrates capable of postural modulation of gas exchange, behavioral closure should systematically inflate TPC-based thermal limit estimates, and the magnitude of this confound should scale with the degree of postural modulation across taxa. We test this prediction in an undescribed sea anemone within the genus *Urticina* (Actiniaria: Actiniidae) distributed along the Northwest Atlantic coast from Newfoundland to Maine. Because no monitoring program targets this undescribed anemone, its distribution must be inferred from opportunistic records. We use iNaturalist observations and correct their spatial and taxonomic biases with a co-occurring effort proxy before modeling distribution. We integrate three complementary methods: (1) a negative binomial regression of effort-corrected encounter rate against environmental predictors spanning both marine (e.g., sea surface temperature, chlorophyll, sea ice) and atmospheric variables (e.g., cloud cover, wind speed), whose retained predictors screen the covariate set for (2) a maximum entropy species distribution model built from the same effort-corrected iNaturalist data, and (3) experimental thermal performance curves from closed-chamber respirometry. We quantify the behavioral confound and develop a multiplicative model that separates behavioral from physiological components of the metabolic response, then assess how the choice of critical thermal maximum (*CT*_max_) estimation method affects thermal safety margin calculations.

## Materials and Methods

### Study organism

The focal animal is an undescribed species of *Urticina*, placed in the genus on the basis of gross morphology (L. G. Harris, personal communication). It is a small anemone, with a pedal disk typically 25 mm in diameter and rarely exceeding 50 mm, attached to hard substrata in the intertidal zone and most abundant at mid-shore, often at the base of fucoid algae or within shaded tidepools on wave-exposed or high-current shores. The column is smooth and rounded and bears no verrucae; unlike *Urticina felina* (Sanamyan *et al*. 2020), it does not retain gravel or shell fragments. Color varies from uniformly red through red-and-beige mottling to entirely beige or brown, with individuals at a given shore tending toward a single hue. The oral disk is translucent and usually matches the column, with the mesenteries visible through it and the tentacle bases occasionally pigmented. Roughly 60 tentacles are arranged in four cycles, each slightly longer than half the oral-disk width, with rounded to faintly clavate tips, and an oral disk that is frequently and uniformly banded (Figure 1).

**Figure 1.**
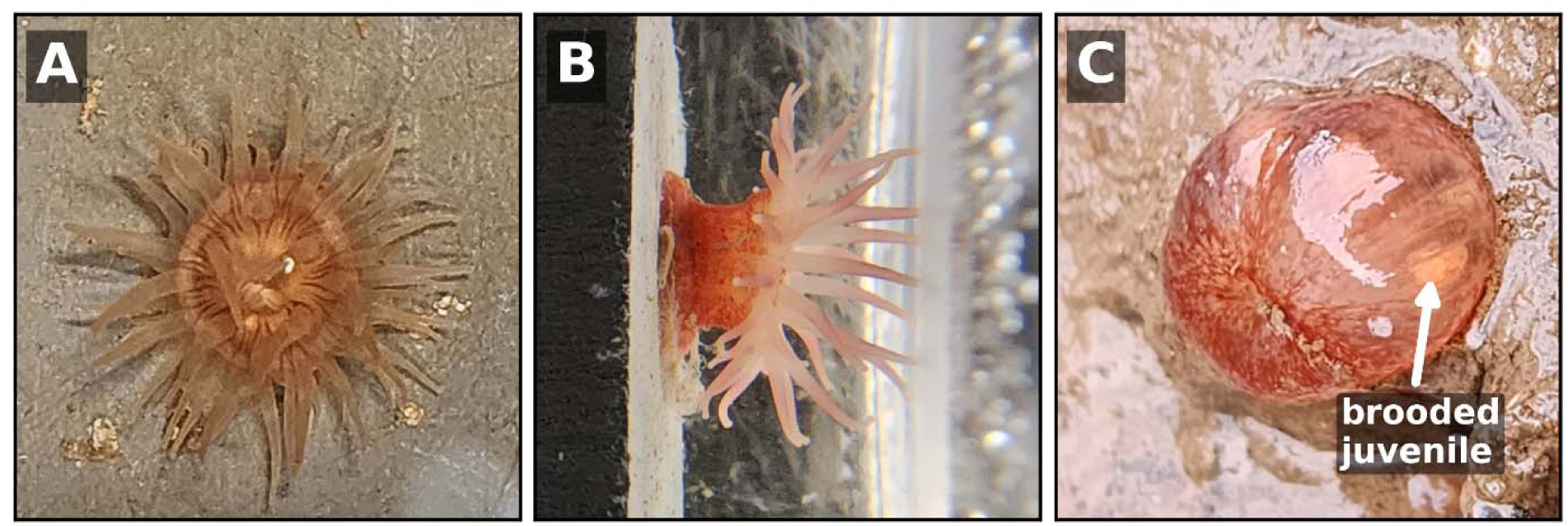
The undescribed *Urticina* sp. (A) Expanded individual in top view, showing the oral disk and the banding that is often present. (B) Side view of an individual attached to an experimental tile, showing the smooth column and tentacle crown. (C) A contracted individual; the column is smooth and lacks verrucae and adherent debris, and a brooded juvenile (orange-beige) is visible within the gastrovascular cavity (arrow).

The species is most readily confused with *Urticina crassicornis*, from which it is distinguished by its smaller size, consistently intertidal distribution, and reproduction at sizes that would be considered juveniles of *U. crassicornis*. *U. crassicornis* occurs subtidally in the region, emerging into the low intertidal only on the largest tides.

Individuals brood young from when pedal disk diameter exceeds 20 mm, and these brooded juveniles can be visible within the gastrovascular cavity (Figure 1C). They are preyed upon by the nudibranch *Aeolidia papillosa* and the sea spider *Pycnogonum littorale*. These field characters distinguish the species *in situ* from its regional congeners, and every occurrence record analyzed here is supported by a photograph archived on iNaturalist (Figure 2).

**Figure 2.**
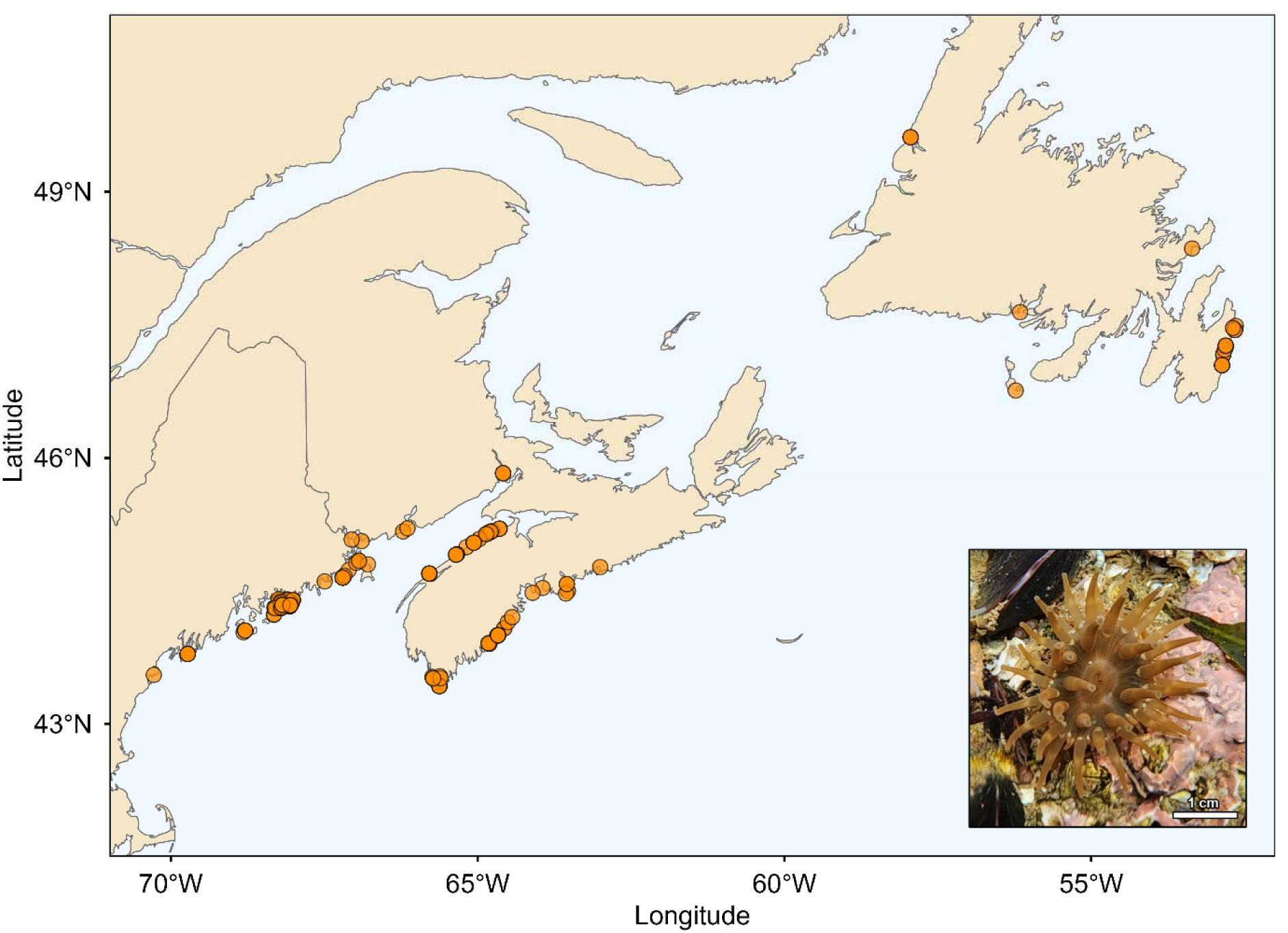
Study area spanning the Northwest Atlantic coast from Massachusetts to Newfoundland. Points indicate iNaturalist observations of the undescribed *Urticina* sp. (bottom right inset).

### Replication statement

### Study area and species occurrence data

All iNaturalist observations of the undescribed *Urticina* within the study extent were obtained on 12 July 2026 (41.5-51°N, 71-52°W, Figure 2) by examining all Actiniaria observations, yielding 357 records. The study extent encompassed the coasts of Massachusetts, New Hampshire, and Maine (USA); New Brunswick, Quebec, Nova Scotia, Prince Edward Island, and Newfoundland and Labrador (Canada); and Saint Pierre and Miquelon (France).

To standardize the spatially variable observation effort found in iNaturalist, a hexagonal grid (H3 resolution 5, ∼250 km^2^, O’Brien 2023) was constructed using all cells that contained shoreline within the extent. Shoreline presence was determined by measuring the length of 1:10m Natural Earth coastline (R package rnaturalearth 1.2.0, Massicotte & South 2025) passing through each cell; cells with zero shoreline length were excluded. Because iNaturalist coordinates carry spatial uncertainty from imprecise georeferencing and observer-obscured locations, some *Urticina* observations fell in cells that did not contain both land and ocean; counts in these cells were reassigned to the nearest coastal cell by centroid distance, as this species is strictly intertidal.

In addition, within each cell, the number of observations of non-target marine species was quantified as an estimate of sampling effort. All observations were filtered within iNaturalist to the animal groups Echinodermata (urchins and sea stars), Crustacea (crabs, shrimp, and isopods), Caenogastropoda (most marine snails), and Thecostraca (barnacles), and the algal groups Rhodophyta (red algae), Ulvophyceae (green algae), and Phaeophyceae (brown algae). Within these groups, there are several taxa that live in freshwater, terrestrial, or pelagic habitats and are therefore not useful for estimating intertidal sampling effort; these groups were removed. They were the Viviparidae (river snails), Oniscidae (wood lice), Asellidae (freshwater isopods), Branchiopoda (branchiopods), Copepoda (copepods), Astacoidea (freshwater crayfish), and Trentepohliales (orange terrestrial algae).

### Environmental predictors

Because this species inhabits the air-water interface, environmental predictors were drawn from both marine and terrestrial atmospheric sources spanning sea surface and benthic variables, atmospheric analyses, terrestrial climate, anthropogenic disturbance, and protected-area coverage (sources, resolutions, and citations in Table S1). These axes were chosen to capture the environmental pressures most likely to bound a cold-water intertidal anemone: thermal exposure during immersion and aerial emersion, winter ice disturbance, planktonic food supply, and anthropogenic habitat modification.

All raster values were extracted as the mean across each cell. For land-dominated cells where marine rasters contained no ocean pixels, values were filled from the mean of immediate neighbors within a 1-ring adjacency; cells remaining without values after this step (more than one cell deep into land with no ocean-adjacent neighbors) were excluded from modeling. Collinearity among the 16 candidate predictors was assessed using variance inflation factors to ensure that each predictor carried independent information. Variables were iteratively removed starting with the highest variance inflation factor until all remaining predictors fell below a threshold of 5.0.

### Encounter rate regression

To assess which environmental predictors structure *Urticina* encounter rate, a negative binomial regression (NB) was fit (R package glmmTMB 1.1.10, Brooks *et al*. 2017) in R 4.6.0 (R Core Team 2026) with observation count as the response, log-transformed effort as an offset, and all variance-inflation-factor screened environmental predictors as covariates. A zero-inflated negative binomial was attempted first but the zero-inflation submodel exhibited complete separation; a DHARMa diagnostic (R package DHARMa 0.4.7, Hartig 2024) confirmed that the standard NB’s dispersion parameter and log-effort offset adequately captured the excess zeros, so the zero-inflation component was dropped.

### Species distribution model

Predictors with p <0.10 in the encounter rate regression were carried forward into a maximum entropy (MaxEnt) species distribution model (Guisan & Zimmermann 2000). This screening threshold is more inclusive than the α = 0.05 convention because the goal is variable reduction and selection rather than hypothesis testing, and the count-based regression’s larger effective sample size retains more information to detect ecologically relevant associations than the binary presence-absence framework of the species distribution model (Aarts, Fieberg & Matthiopoulos 2012). The MaxEnt model was fit at the same hex-level as the negative binomial regression, with model tuning and spatial block cross-validation (R packages maxnet 0.1.4 and ENMeval 2.0.5.2, Kass *et al*. 2021; Phillips 2021). Since predictors were extracted as cell-level zonal means, presences were thinned to one record per cell. Background was sampled with replacement from the coastal cells weighted by effort to correct for spatially heterogeneous citizen-science detection probability (Phillips *et al*. 2009; Fourcade *et al*. 2014).

MaxEnt models can range from simple linear responses to complex nonlinear interactions, and overly complex models risk fitting noise rather than ecological signal (Radosavljevic & Anderson 2014). To guard against this, model complexity was tuned by evaluating 12 candidate models spanning two levels of response shape (linear; linear and quadratic) crossed with six levels of regularization strength (0.5, 1.0, 1.5, 2.0, 3.0, and 5.0), where higher regularization penalizes complexity more heavily. Models were evaluated using spatial block cross-validation (R package blockCV 3.2.0, Valavi *et al*. 2019), which partitions the study area into geographically distinct folds to prevent spatially autocorrelated training and test data from inflating performance estimates. The optimal model was selected by maximum cross-validated AUC. Where validation AUC was tied, we retained the higher regularization multiplier as the more conservative choice at this sample size, though no tie arose in the final tuning grid; higher-than-default regularization reduces overfitting in MaxEnt models generally (Radosavljevic & Anderson 2014).

### Thermal performance experiment

To assess whether standard respirometric methods reliably characterize thermal limits in this species, and to test for behavioral confounds in the measured metabolic response, we measured thermal performance curves experimentally. *Urticina* sp. were collected from West Quoddy Head, Lubec, ME (44.817°N, 66.950°W) by hand from the mid-intertidal and transported to the Schiller Coastal Studies Center, Harpswell, ME. There, they were maintained in 25-µm filtered seawater in flow-through sea tables, fed a diet of blue mussel (*Mytilus edulis*) every two to three days, and held at ambient temperature under natural lighting.

Anemones were placed in custom sealable acrylic respirometry chambers atop VWR 230 submersible stirrers (speed 3), with perforated plastic dividers separating the magnetic fleas from the anemones to allow circulation while preventing contact. The chambers, initially open to make certain oxygen was fully saturated, sat in a temperature-controlled flow-through water bath supplied with 0.2-µm filtered and UV-treated seawater from Harpswell Sound; an airstone in the bath maintained 100% oxygen saturation. Bath temperature was controlled by two 500 W Inkbird ITC-308-WIFI heaters and two ARCTICA DBM-250 titanium chillers plumbed in sequence.

Mass-specific oxygen consumption rates (µg O_2_ g^-1^ h^-1^) were measured for 18 individual *Urticina* sp. across seven temperature treatments (1, 5, 10, 15, 20, 25, 30°C; n = 126 total trials) using closed-chamber respirometry. Anemones were tested at sequentially warmer temperatures starting at 1°C, with at least 48 h between trials and 72 h if fed to prevent digestion-related metabolic elevation. This sequential design held individual identity constant across temperatures, removing the large between-individual variation in baseline metabolic rate from the temperature contrast; randomizing temperature order would have risked premature mortality at the warmest treatment. Each trial began with a 15-min acclimation after which chambers were sealed. Closed-chamber measurement was chosen because the absolute oxygen uptake of animals this size is resolvable only over multi-hour incubations, and pilot studies informed trial durations to keep chamber oxygen above respiration-limiting levels as suggested by Killen *et al*. (2021); oxygen did not fall below 68% of the initial concentration in any trial (Table S2). Oxygen was measured at the start and end of each trial with a Strathkelvin Instruments oxygen meter (Model 872) and electrode, calibrated daily with an anoxic sodium sulfite solution and a 100% oxygen-saturated water sample; a YSI ProDSS provided water temperature, salinity, and pressure for electrode calibration. Consumption rates were control-corrected and normalized to dry mass. Dry mass was measured by drying each anemone at the conclusion of the experiments on a pre-weighed weigh boat at 60°C for seven days and weighing to the nearest 0.1 mg. Anemone expansion state was scored from 0 (fully closed) to 1 (fully expanded) to the nearest 0.2 by a single observer, T.M. This was recorded at the start, at one-quarter, one-half, and three-quarters of the way through each trial, and immediately before the end; the mean of these five observations was the behavioral covariate. Animals began the acclimation period expanded.

To jointly estimate the thermal response and the confounding effect of behavioral state, Bayesian nonlinear mixed-effects models were fit to the full individual-level dataset (R package brms 2.22.0, Bürkner 2017). Because the choice of functional form can itself determine the inferred thermal limit, symmetric and asymmetric shapes were compared rather than assuming one form (Angilletta 2006). Four TPC functional forms were crossed with three behavioral structures, producing 12 candidate models. The four TPC shapes were Gaussian, quadratic, Briere2 (Brière *et al*. 1999), and Sharpe-Schoolfield with high-temperature inactivation (Schoolfield, Sharpe & Magnuson 1981). The three behavioral structures formalized competing hypotheses about how expansion state affects metabolic rate: (1) naïve (behavioral state ignored), (2) additive (expansion state contributes a temperature-independent offset), and (3) multiplicative, in which expansion state proportionally scales the entire TPC:

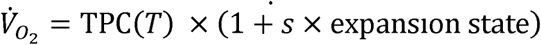

Under the multiplicative formulation, a fully closed anemone (expansion state = 0) expresses the baseline TPC, while a fully expanded anemone expresses (1 + *s*) times the baseline, where *s* is estimated from the data. This parameterization reduces the identifiability problems that arise when TPC parameters and behavioral effects are estimated independently in a two-stage approach. This formulation assumes linear scaling between expansion state and metabolic rate.

All models included a random intercept for individual identity on the amplitude parameter to account for repeated measures across temperatures. Weakly informative normal priors were specified for all parameters, centered on plausible physiological values (Table S3). Models were fit with four chains of 4000 iterations (2000 warmup) and convergence was assessed via the Gelman-Rubin diagnostic (R < 1.05) and fewer than 20 divergent transitions across the 8000 post-warmup samples. Chains that met these thresholds on the first attempt were retained; chains that failed were refit with up to four alternative random-number seeds; if all four still failed, the model was considered non-convergent. Three additional TPC shapes (Lactin2, Thomas 2012, Beta 2012) were excluded from the main comparison after prototype fits failed to converge (Table S4, models are described in Lactin *et al*. 1995; Niehaus *et al*. 2012; Thomas *et al*. 2012). Converging models were compared using leave-one-out cross-validation information criterion (LOOIC, R package loo 2.8.0, Vehtari, Gelman & Gabry 2017), which estimates out-of-sample predictive performance while accounting for the effective number of parameters. TPC parameters (*T_opt_*, *R_max_*) and behavioral multiplier (*s*) were summarized as posterior medians with 95% credible intervals. Because symmetric TPC models can extrapolate *CT_max_*far beyond the observed temperature range, *CT_max_* was derived numerically as the temperature above *T_opt_* at which the predicted rate drops below 5% of *R_max_*. For the Briere2 model, this closely corresponds to its explicit *t*_max_ parameter where rate = 0. *CT_max_* is also reported from direct experimental observation.

To assess whether the inferred thermal parameters were robust to among-individual variation in body size, we re-fit the best-performing TPC across all three behavioral structures with the centered logarithm of dry mass added as a fixed effect on the model’s amplitude parameter. These mass-adjusted variants used the same priors and behavioral structures as the corresponding base models, with a stricter target acceptance probability of 0.99 and maximum tree depth of 13 to handle the tighter posterior geometry. Pairwise comparison of the mass-adjusted and unadjusted variants by LOOIC tested whether explicitly modeling body size altered the inferred *T_opt_*, *R_max_*, or behavioral offset or multiplier.

## Results

### Encounter rate regression

Environmental predictors structured *Urticina* encounter rate. Collinearity screening dropped pH, air temperature, vapor pressure deficit, and solar radiation. *Urticina* was present in 41 of 672 cells with complete predictor data; the negative binomial regression converged (dispersion θ = 0.152; Table S5) with three predictors significantly associated with *Urticina* encounter rate. Cloud cover was the strongest positive predictor (IRR = 5.49 per SD; 95% CI: 1.97-15.28; *p* = 0.001), percent developed coastline was the strongest negative predictor (IRR = 0.51 per SD; 95% CI: 0.34-0.77; *p* = 0.002), and warmer winter minimum SST was associated with higher encounter rate (IRR = 2.70 per SD; 95% CI: 1.06-6.90; *p* = 0.038). An additional predictor fell between p = 0.05 and the p = 0.10 screening threshold for inclusion in the MaxEnt model: chlorophyll-a (borderline at p = 0.051; IRR = 2.40 per SD; 95% CI: 1.00-5.78). Predicted encounter rate was highest along the coasts of Maine, Nova Scotia, and southern Newfoundland, and dropped toward both the northern limit in Newfoundland and the southern limit in mid-Maine (Figure 3A).

**Figure 3.**
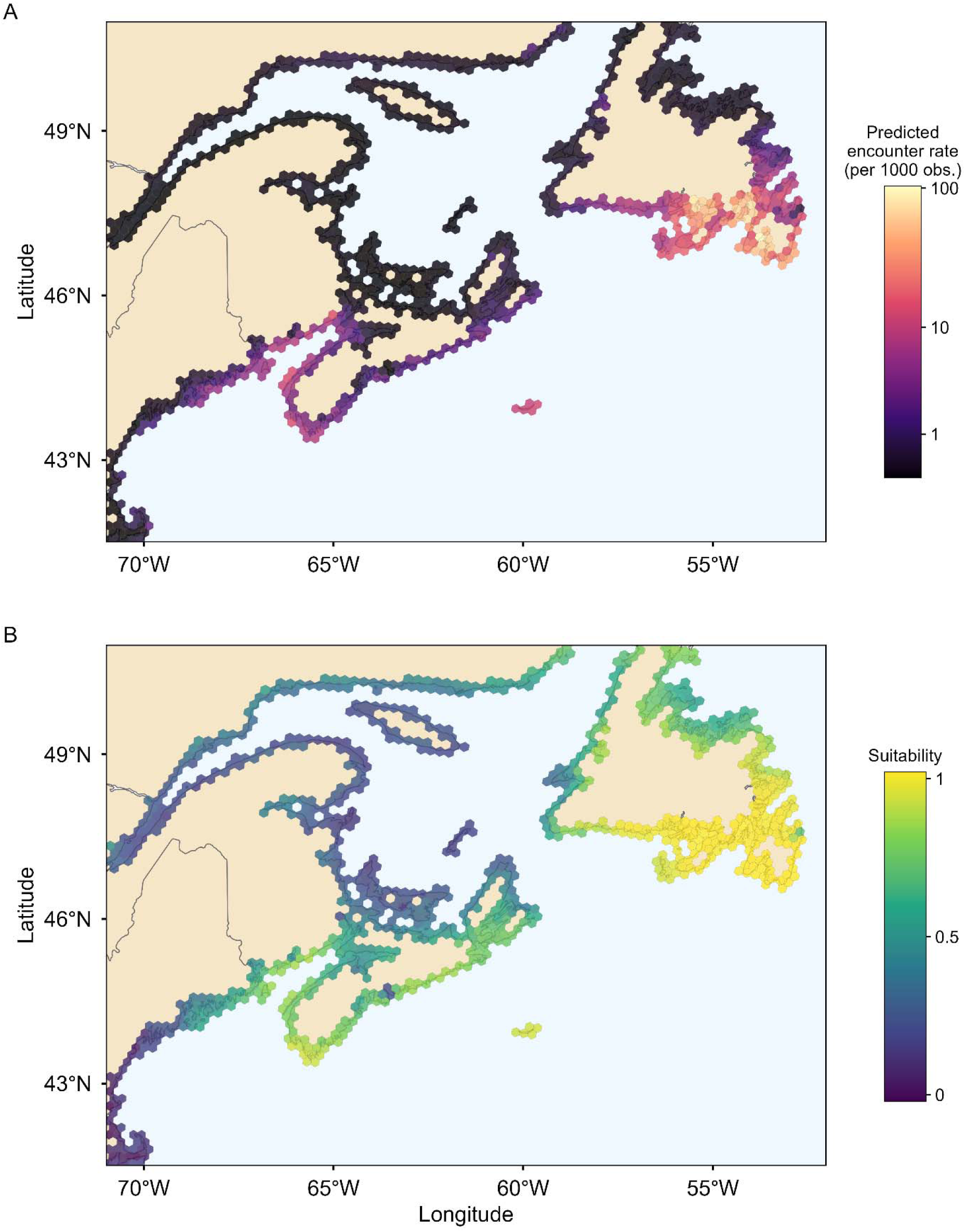
Comparison of (A) negative binomial predicted encounter rate and (B) MaxEnt habitat suitability. Encounter rate is the expected *Urticina* sp. count per 1000 iNaturalist observations, predicted from the negative binomial regression at median effort. Both panels show all coastal cells with complete environmental predictor data.

### Species distribution model

The MaxEnt model used four NB-screened predictors, 41 spatially thinned presence cells, and 5000 effort-weighted background draws from 848 coastal cells with environmental data. The number of cells is higher because the negative binomial requires a nonzero effort offset, so cells without effort drop out, whereas the MaxEnt model does not. The optimal configuration used linear and quadratic feature classes with a regularization multiplier of 5.0 (validation AUC = 0.861 ± 0.073; training AUC = 0.842; AUC gap = -0.019). Of the four candidate predictors, three contributed non-zero features (Figure S1), led by winter minimum SST, followed by cloud cover and developed coastline. Predicted habitat suitability was highest along the coasts of Maine, Nova Scotia, and southern Newfoundland, and dropped toward both the northern detection limit in Newfoundland and the southern limit in mid-Maine (Figure 3B). The two models agreed on where *Urticina* should be common: the geographic pattern in MaxEnt suitability tracked NB-predicted encounter rate across the same coastal cells with both surfaces peaking in the Gulf of Maine and the Bay of Fundy and falling off at both range edges (Spearman ρ = 0.82; Figure 4).

**Figure 4.**
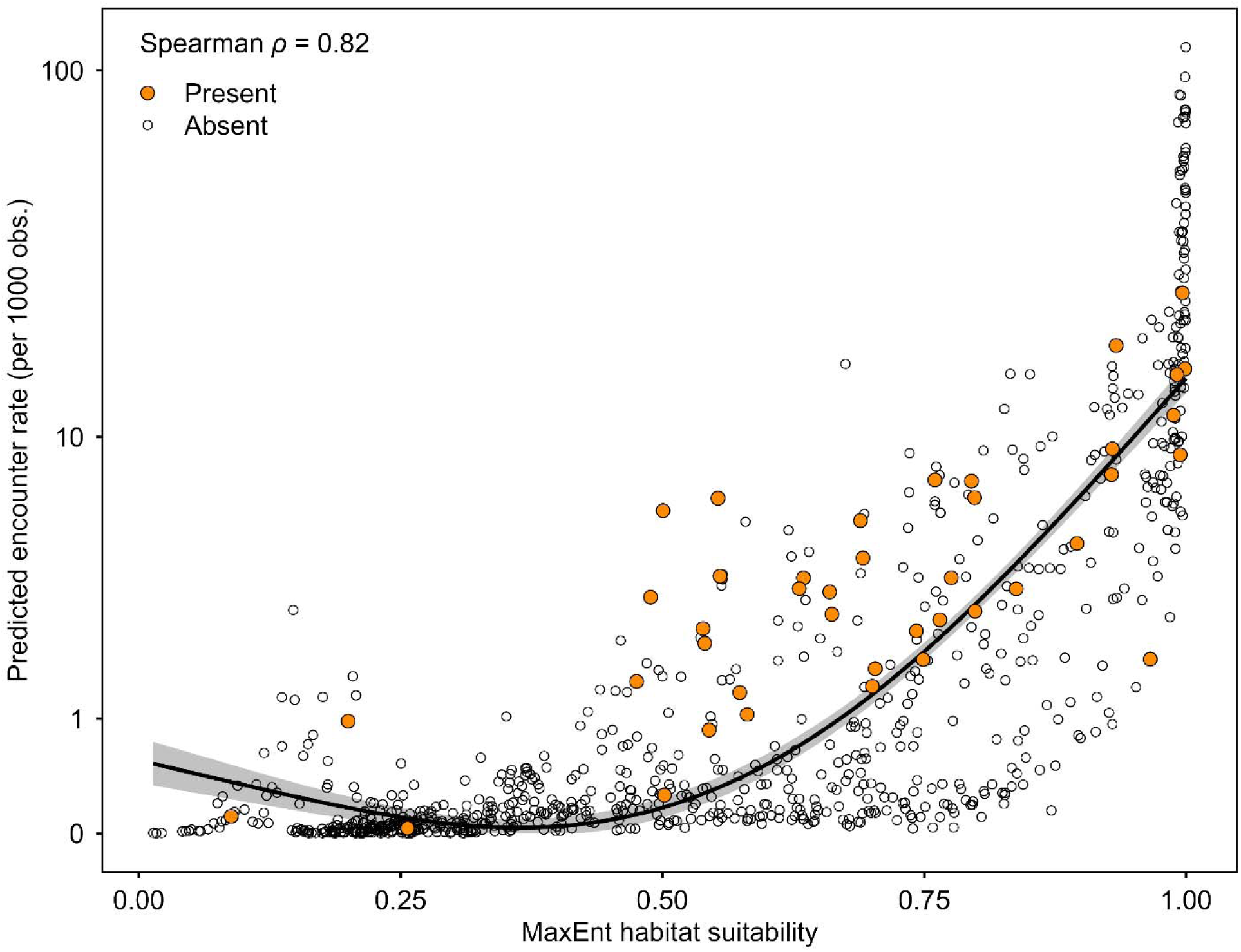
Agreement between negative binomial predicted encounter rate (per 1000 iNaturalist observations at median effort) and MaxEnt habitat suitability across the same coastal cells. Orange points mark cells with at least one confirmed *Urticina* sp. observation; open points mark cells with no record. The black line is a GAM smooth (k = 3) fit to all cells; the gray band is the 95% confidence interval.

### Thermal performance experiment

Mean mass-specific oxygen consumption rose from 125 µg O_2_ g^-1^ h^-1^ at 1°C to a peak of 335 µg O_2_ g^-1^ h^-1^ at 25°C, then fell to 236 µg O_2_ g^-1^ h^-1^ at 30°C (Figure 5). All 18 individuals died at 30°C, placing the empirical lethal limit between 25 and 30°C. Expansion state was partial and variable across the coldest treatments (means 0.26-0.51 from 1 to 20°C) but declined above 20°C, from 0.43 at 20°C to 0.26 at 25°C, before collapsing to 0.05 at the lethal 30°C treatment.

**Figure 5.**
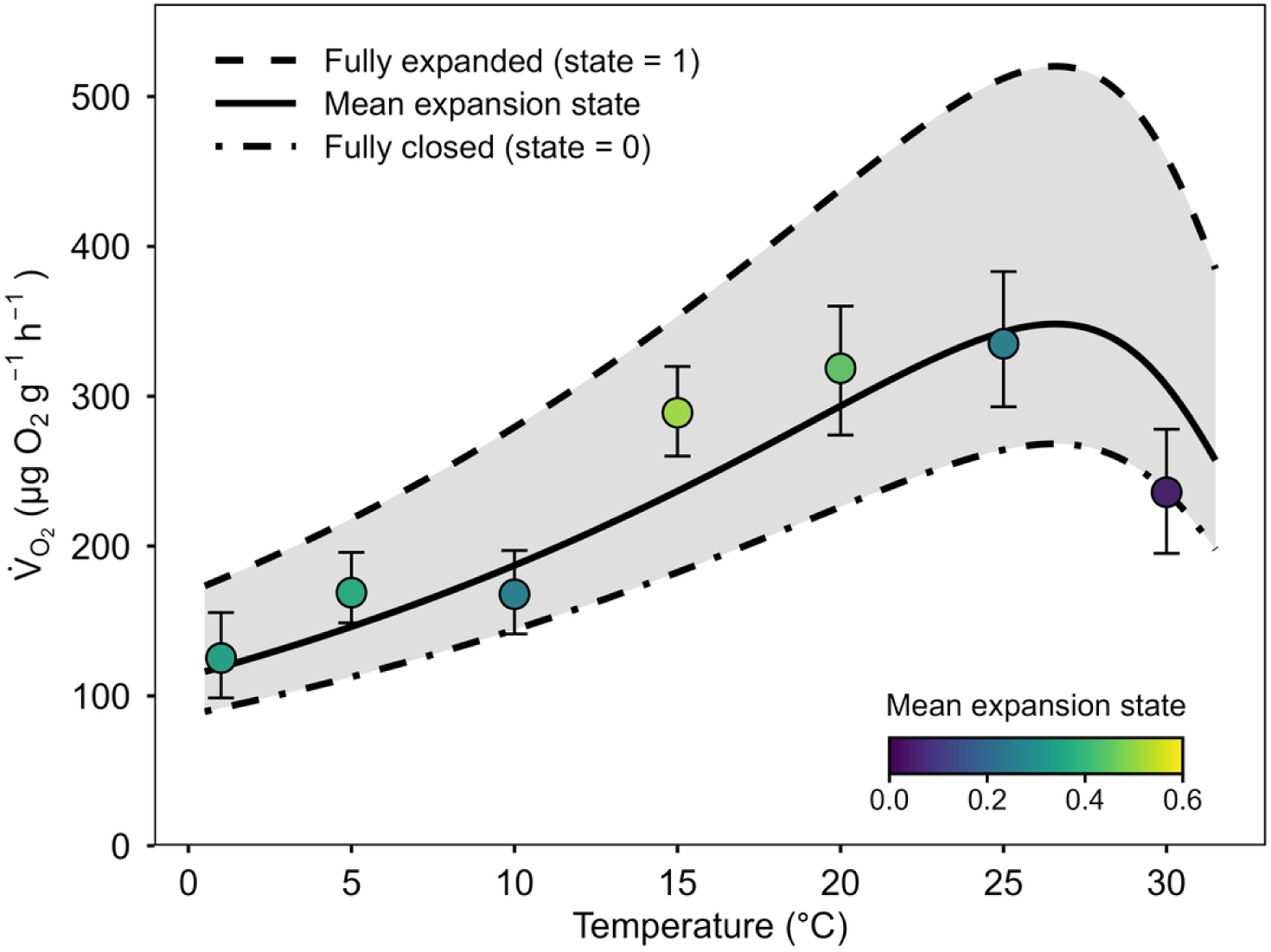
Thermal performance of *Urticina* sp. under the best thermal performance model. The solid line is the model prediction at the mean observed expansion state (0.32); the dashed and dot-dash lines are the predictions at fully expanded (state = 1) and fully closed (state = 0), with the shaded region between them spanning the rescaling that the multiplicative behavioral term applies across temperature. Points and error bars show per-temperature means and 95% bootstrap confidence intervals of the measured mass-specific oxygen consumption with point color indicating the mean observed expansion state at each temperature.

All 12 candidate models converged (Table 2). Every behavioral structure improved fit over the naïve formulation with the smallest ΔLOOIC of 16.8 (SE = 5.3) between the Briere2 naïve and multiplicative formulations. The Sharpe-Schoolfield multiplicative model was the best performing model, estimating a thermal optimum of 26.5°C (95% CI: 24.7-28.1) and a peak metabolic rate of 349 µg O_2_ g^-1^ h^-1^ (95% CI: 311-394). Critical thermal maximum from this model was 45.9°C (95% CI: 35.3-65.4; Table 2), compared with 74.2°C from the symmetric Gaussian multiplicative fit. Briere2 was the only shape for which the additive structure outperformed the multiplicative (ΔLOOIC = 46.7).

**Table 1.** Replication statement for experimental and observational components of this study.

| Scale of inference | Scale at which the factor of interest is applied | Number of replicates at the appropriate scale |
| --- | --- | --- |
| Species | Hexagonal grid cell (~250 km) | 672 (encounter rate) and 848 cells (suitability), 41 with anemones |
| Species | Individual anemone | 18 from one site, 7 temperatures each |

**Table 2.** Model performance comparison of Bayesian thermal performance models.

| TPC shape | Behavioral structure | $T_{\text{opt}}$ (°C) | $CT_{\text{max}}$ (°C) | $\Delta\text{LOOIC}$ |
| --- | --- | --- | --- | --- |
| Sharpe-Schoolfield | Multiplicative | 26.5 | 45.9 | 0.0 |
| Sharpe-Schoolfield | Additive | 25.4 | 45.9 | 2.2 |
| Gaussian | Multiplicative | 28.8 | 74.2 | 5.1 |
| Gaussian | Additive | 24.5 | 57.0 | 11.7 |
| Quadratic | Multiplicative | 39.5 | 85.0 | 18.9 |
| Briere2 | Additive | 24.9 | 39.1 | 23.8 |
| Quadratic | Additive | 27.9 | 64.8 | 34.5 |
| Sharpe-Schoolfield | Naïve | 23.8 | 44.8 | 42.0 |
| Gaussian | Naïve | 21.9 | 54.7 | 55.8 |
| Briere2 | Multiplicative | 23.7 | 39.0 | 70.5 |
| Quadratic | Naïve | 22.3 | 47.2 | 73.7 |
| Briere2 | Naïve | 22.3 | 37.5 | 87.3 |

Posterior predictive checks and residual diagnostics of the Sharpe-Schoolfield multiplicative model showed no systematic departures (Figure S2, Figure S3). Under the multiplicative structure, a fully expanded anemone respired 1.94 times the rate of a fully closed anemone at any temperature (behavioral multiplier *s* = 0.94, 95% CI: 0.63-1.30; Figure 5). Across the observed range of expansion (0.05 to 0.51) the difference was about 1.4-fold; the 1.94-fold value compares the model’s fully closed and fully open states, which were not directly observed. Excluding the lethal 30°C treatment left the behavioral multiplier modestly lower, with overlapping credible intervals (*s* = 0.82, 95% CI 0.57-1.10), confirming that the behavioral partition does not depend on the single lethal treatment; the thermal optimum, by contrast, is informed by that treatment and should be read as conditional on it. Among-individual variation in baseline metabolic rate was substantial (random-intercept SD = 54 µg O_2_ g^-1^ h^-1^; 95% CI: 5.4-114, Figure S4). This individual-level variation correlated negatively with body size (Spearman ρ = - 0.59, *p* = 0.012; Figure S5); adding the logarithm of dry mass as a fixed effect on the reference-temperature rate improved the Sharpe-Schoolfield multiplicative fit (ΔLOOIC = 6, SE = 2.4) but left the thermal optimum, peak rate, and behavioral multiplier unchanged (Supplementary Table S6).

## Discussion

Standard thermal performance curves overshoot the observed lethal limit in *Urticina* by a wide margin. Every individual died during the 30°C respirometry experiment, yet a symmetric Gaussian fit extrapolates the upper thermal limit to 74.2°C, a value that would place a cold-water North Atlantic anemone among the most heat-tolerant marine invertebrates on Earth. Even the best-fitting asymmetric model, which is explicitly bounded, puts the upper thermal limit at 45.9°C, nearly 16°C above the temperature at which the animals actually died. That curve-fitted maximum is sensitive to the high-temperature inactivation priors and lies well beyond the observed data, so we treat the empirical 25-30°C lethal bracket as the warm-limit metric. Thermal performance is strongly asymmetric about the optimum (Kingsolver 2009), yet symmetric functions such as the Gaussian and quadratic carry no explicit upper-limit parameter (Table S3), so their descending limb is extrapolated from the ascending limb rather than estimated from data near the limit. Information criteria reward fit to the ascending half of the curve without penalizing implausible extrapolation of the descending half. In some animals, that descending half reflects behavioral closure as much as physiology. What standard curve-fitting reads as a gradual physiological decline is actually an animal retracting before it dies. That retraction is likely adaptive in nature, a behavioral attempt to reduce exposure during brief thermal extremes; however, in this experiment the 30°C treatment exceeded the threshold at which closure could rescue the animal.

A single additional parameter recovers the physiological signal. Fitting the measured metabolic rate as a temperature-dependent baseline multiplied by a behavioral term (s) yielded a fully expanded *Urticina* respiring 1.94 times the rate of a fully closed one at the same temperature (s = 0.94, 95% CI: 0.63-1.30). A transition from fully expanded to fully closed therefore suppresses the measured respiration by about 48% independent of any change in physiology. Note that the multiplicative model makes a specific prediction: behavior changes the height of the TPC, not the temperature at which it peaks. Recovering the physiological signal therefore raises the predicted rate at every temperature uniformly, and thermal optimum at fully expanded (state = 1) lies within 0.2°C of the optimum temperature at mean expansion state. Mortality at 30°C reflects the failure of protective closure to keep the animal alive, not a higher physiological optimum hidden by that closure. Standard respirometry folds this suppression into the measured response without separating it, understating the temperature at which genuine physiological impairment begins.

The same pattern appears across other invertebrates: behavioral withdrawal at high temperature suppresses measured respiration before molecular stress markers register (Ortmann & Grieshaber 2003; Anestis *et al*. 2007). The mechanism takes different forms across taxa, including valve closure in bivalves that drops respiration to roughly 10% of standard, anticipatory cardiac depression that produces non-monotonic TPCs in intertidal oysters, and postural shifts that can reverse the rank order of metabolic rates among individuals of a single anemone species (Ortmann & Grieshaber 2003; Hui *et al*. 2020; Maskrey *et al*. 2020; Maskrey *et al*. 2024). A two-process framework that explicitly partitions metabolic depression treats such metabolic depression as a common response rather than an exception (Liao *et al*. 2021). Behavioral and physiological TPCs decouple more generally in intertidal ectotherms (Monaco, McQuaid & Marshall 2017).

In every case, the measured response is a mixture of physiological and behavioral processes that a single-shape TPC cannot separate. Here we treat the measured rate as a physiological baseline scaled by behavior, which separates the two components with one additional parameter and one additional data stream: a quantitative behavioral metric recorded alongside oxygen consumption.

These findings quantify the confound in one species; whether its magnitude scales across taxa with the degree of postural regulation of gas exchange, as we predict, remains to be tested. Marine invertebrate taxa, with the strongest postural modulation of gas exchange (e.g., valve closure in bivalves, contraction in anemones, retraction in tunicates, lophophore withdrawal in bryozoans), should show the largest gap between naïve and behavior-corrected TPCs. The multiplicative model provides a standardized framework for measuring that gap. If the confound operates broadly, it carries concrete implications for climate vulnerability assessment. At the warmest occupied site, where the warmest summer sea surface temperature is 20.9°C, the behavioral safety margin for *Urticina* is roughly 4 to 9°C, the distance to the 25-30°C bracket at which sustained closure sets in, and several factors suggest this margin is an overestimate. Chronic thermal tolerance yields margins roughly half those from acute assays, marine heatwave pre-exposure can substantially reduce upper thermal limits, and ambient sea surface temperature understates operative body temperature during aerial exposure (Helmuth *et al*. 2002; Helmuth *et al*. 2006; King *et al*. 2025; Molina *et al*. 2025). The potentially generous safety margins reported for many marine taxa may therefore reflect unmeasured behavioral suppression rather than real physiological headroom. Because the rate of heat failure escalates nonlinearly above the sublethal stress range (Jørgensen *et al*. 2022), even modest overestimates of thermal tolerance translate into large errors in projected vulnerability.

The distributional data are consistent with a niche structured by temperature and exposure, though they do not by themselves isolate summer heat as the warm-edge driver; the retained predictors fall into three ecologically coherent groups. Variables that buffer thermal stress at the air-water interface carry consistent positive signals: both distribution models flagged higher cloud cover (a proxy for fog and reduced vapor-pressure deficit, which reduce desiccation stress) as important, and *Urticina* were more likely to be found in regions with warmer winters and less sea ice. The winter-warming association, however, is significant only in the encounter-rate model, with a negligible MaxEnt coefficient. Anthropogenic disturbance, captured by percent developed coastline, carries the largest negative effect in both models, consistent with impervious runoff and physical disturbance degrading rocky intertidal habitat at urbanized coasts despite higher sampling effort by citizen scientists in those regions. Because developed coastline peaks in the urbanized south near the warm range edge, its negative effect cannot be fully separated from summer temperature there. Encounter rate and habitat suitability both decline across southern Gulf of Maine temperatures, and experimentally measured thermal performance peaks at 26.5°C, just inside the 25-30°C lethal bracket. The encounter-rate signal flags this warm edge from citizen-science observations, the MaxEnt suitability surface flags it from environmental space, and the chamber experiment localizes it to 25-30°C. These lines converge, but the negative binomial and MaxEnt models share the same occurrence records and screened predictors, so their agreement is a consistency check rather than independent corroboration; warm-edge thermal limitation rests on the experiment.

For this undescribed *Urticina*, a narrow behavioral safety margin combined with the Gulf of Maine’s rapid warming (Pershing *et al*. 2015; Thomas *et al*. 2017) places warm-edge populations within a few degrees of functional thermal failure. Thermal mismatch has already driven the failure of at least one sea anemone in this region (Wells 2013; Wells & Harris 2019), and metabolic depression under combined thermal and acidification stress is documented in Gulf of Maine *Mytilus edulis* from the same habitat (Lesser 2016). Temperature-oxygen interactions and ocean acidification may narrow the effective thermal window further through pathways single-stressor respirometry cannot detect. American lobster (*Homarus americanus*) has shown a parallel pattern: as the southern New England stock collapsed under warming-driven recruitment failure while the Gulf of Maine fishery surged on the same temperature trend, the recruitment optimum tracked northeastward into Maine and Atlantic Canada (Pinsky *et al*. 2013; Le Bris *et al*. 2018). Whether *Urticina* shifts poleward depends on factors a static niche model cannot capture, namely access to local thermal refugia, larval dispersal, and acclimation capacity (Chapperon & Seuront 2011; Bates *et al*. 2014). Its undescribed taxonomic status compounds the conservation risk: undescribed species are not usually included in formal assessments (Liu *et al*. 2022).

More broadly, standard respirometric TPCs can produce misleading thermal limit estimates for sessile organisms when behavioral state is not measured, and that a single additional parameter can recover much of the physiological signal. The immediate priority is to test whether the confound scales across sessile taxa as predicted; applying the multiplicative model to shelled mollusks, tunicates, and other sea anemones would determine whether the behavioral contribution documented here generalizes. Until such data exist, safety margins derived from standard TPCs for sessile marine invertebrates should be treated as upper bounds. Citizen science platforms, which now provide distributional baselines, offer a practical way to extend these thermal-risk assessments to the many intertidal species that still lack formal thermal physiology data.

## Acknowledgements

We thank the iNaturalist community for contributing the biodiversity observations that made the distributional component of this study possible. Thank you to Barry Logan and Amy Johnson for reading early versions of the manuscript. Thank you to the staff and visiting researchers at the Bowdoin College Schiller Coastal Studies Center and Biology Department for your help with laboratory assistance and logistical support. Special appreciation for the efforts and company of Jonathan Allen, Heidi Franklin, Holly Parker, Jaret Reblin, and Rachel Reuling. Animals were collected under the permit Maine Department of Marine Resources permit SL2024-11-04. Research involving invertebrate animals at Bowdoin College does not require Institutional Animal Care and Use Committee review under current institutional policy.

## Conflict of Interest

The authors declare no conflict of interest.

## Authors’ Contributions

All authors conceived the ideas, designed methodology, and collected the data; Christopher D. Wells analyzed the data and led the writing of the manuscript. All authors contributed critically to the drafts and gave final approval for publication.

## Data Availability Statement

Data and analysis code available from Zenodo: https://doi.org/10.5281/zenodo.21346544 (Edgar *et al*. 2026). The iNaturalist observations are publicly available at https://inaturalist.org.

## Funding

Funding was provided by the Henry L. and Grace Doherty Charitable Foundation Coastal Studies Research Fellowship and the Doherty Marine Biology Postdoctoral Fund.

## Notes

https://github.com/christopherwells/urticina-thermal-niche

https://doi.org/10.5281/zenodo.21346544

### Competing Interest Statement

The authors have declared no competing interest.

### Summary of Updates

We corrected the units on all oxygen consumption rates. They were labeled in milligrams when the correct unit is micrograms, a factor of one thousand. Only the label was wrong. The measurements, the model fits, and every conclusion are unchanged. We regenerated the figures whose axis labels carried the wrong unit. We corrected the CHELSA citation. The earlier version cited a daily forcing dataset that we never used. We used the CHELSA version 2.1 monthly climatologies, and we now cite both the method paper and the versioned data record. We also corrected the temporal coverage listed for these data to 1981 through 2010. We added reference entries for data sources and thermal performance models that had been cited but were missing from the bibliography. We corrected the reported total number of iNaturalist observations to match the cited source, and we now state the period for the reported warming rate. We added a French translation of the abstract.

