## Supplemental Tables and Figures for "Behavioral metabolic suppression confounds thermal performance estimates and climate vulnerability assessments in a marine ectotherm"

Table S1. Environmental datasets used in this study.

| Dataset | Variables | Resolution | Temporal coverage | Reference |
| --- | --- | --- | --- | --- |
| Bio-ORACLE v3.0 | All water temperatures  Salinity  Chlorophyll-a  pH  Sea ice concentration | 0.05° (~5 km) | 2010-2020 | Assis *et al.* (2024)  Tyberghein *et al.* (2012) |
| ERA5 reanalysis | Cloud cover  Wind speed | 0.25° (~28 km) | Jun-Aug 2000-2020 | Hersbach *et al.* (2023) |
| CHELSA v2.1 | Air temperature  Precipitation  Vapor pressure deficit | 30 arcsec (~1 km) | 1981-2010 | Karger *et al.* (2017)  Karger *et al.* (2021) |
| WorldClim 2.1 | Solar radiation | 5 arcmin (~10 km) | 1970-2000 | Fick and Hijmans (2017) |
| GHSL R2023A | Built-up surface fraction | 10 m | 2020 | Pesaresi and Politis (2026) |
| WDPA | Protected area fraction | Vector polygons | 2026 | UNEP-WCMC and IUCN (2026) |
| Natural Earth 1:10m | Shoreline length | 1:10,000,000 vector | - | Massicotte and South (2025) |

Table S2: Respirometry trial durations by temperature treatment. Durations were determined from pilot studies to keep chamber oxygen above the level that would limit respiration throughout each trial. Higher temperatures required shorter trials due to elevated metabolic rates.

| Temperature (°C) | Mean (h) | SD (min) | Minimum (h) | Maximum (h) |
| --- | --- | --- | --- | --- |
| 1 | 4.1 | 25 | 3.2 | 4.6 |
| 5 | 4.2 | 19 | 3.9 | 4.8 |
| 10 | 4.2 | 6 | 4.0 | 4.4 |
| 15 | 4.1 | 7 | 4.0 | 4.4 |
| 20 | 3.1 | 8 | 3.0 | 3.4 |
| 25 | 2.8 | 18 | 2.3 | 3.3 |
| 30 | 2.0 | 10 | 1.7 | 2.2 |

Table S3: Prior distributions for all TPC model parameters. All priors are normal distributions with the listed mean and standard deviation. Bounds indicate hard parameter constraints enforced during sampling (lb = lower bound, ub = upper bound).

| Parameter | Description | Mean | SD | Bounds | TPC shape(s) |
| --- | --- | --- | --- | --- | --- |
| *R_max_* | Peak metabolic rate | 50 | 20 | lb = 0 | Gaussian |
| *T_opt_* | Thermal optimum | 20 | 10 | - | Gaussian |
| *a* | Curve width (SD) | 15 | 10 | lb = 1 | Gaussian |
| *q_a_* | Intercept | 0 | 30 | - | Quadratic |
| *q_b_* | Linear coefficient | 3 | 3 | - | Quadratic |
| *q_c_* | Quadratic coefficient | -0.05 | 0.1 | ub = 0 | Quadratic |
| *b_a_* | Rate scaling constant | 0.001 | 0.005 | lb = 0 | Briere2 |
| *T_min_* | Lower thermal limit | -2 | 5 | ub = 5 | Briere2 |
| *T_max_* | Upper thermal limit | 32 | 3 | lb = 25, ub = 40 | Briere2 |
| *b_b_* | Shape exponent | 2 | 1 | lb = 0.5, ub = 5 | Briere2 |
| *R_ref_* | Rate at reference temperature | 40 | 20 | lb = 0 | Sharpe-Schoolfield |
| *E_a_* | Activation energy | 0.5 | 0.3 | lb = 0 | Sharpe-Schoolfield |
| *E_ah_* | High-temp inactivation energy | 3 | 2 | lb = 0 | Sharpe-Schoolfield |
| *T_h_^-1^* | Inverse inactivation temp | 0.0033 | 0.0002 | - | Sharpe-Schoolfield |
| *T_ref_^-1^* | Inverse reference temp | 0.00345 | 0.0005 | - | Sharpe-Schoolfield |
| *b_open_* | Additive behavioral offset | 20 | 15 | - | All (additive) |
| *s_1_* | Multiplicative behavioral scaling | 1 | 1 | lb = 0 | All (multiplicative) |

Table S4: Thermal performance curve models excluded from the candidate set because their mathematical formulations could not be reliably sampled. All model combinations were attempted with four independent sampling runs; those that produced inconsistent results across runs (Gelman-Rubin *R̂* > 1.05) or encountered numerical failures during sampling were excluded.

| TPC shape | Behavioral structure | Failure reason |
| --- | --- | --- |
| Lactin2 | All three | Numerical instability: exponential terms caused sampling to diverge regardless of starting values |
| Thomas 2012 | All three | Formula intractable: exponential growth term could not be constrained by the data, producing implausible parameter values |
| Beta 2012 | All three | Failed before sampling began: power terms produced undefined values during initialization |

Table S5. Negative binomial regression coefficients for *Urticina* encounter rate. All predictors are standardized (z-scored); estimates are on the log-count scale. Incidence rate ratios (IRR) give the multiplicative change in expected count per one standard deviation increase in the predictor. The SD column shows the standard deviation of each predictor in its original units.

| Term | Estimate | SE | IRR | SD | *z* | *p* |
| --- | --- | --- | --- | --- | --- | --- |
| Cloud cover | 1.703 | 0.522 | 5.49 | 0.050 prop. | 3.26 | 0.001 |
| Developed surface | -0.673 | 0.213 | 0.51 | 0.008 prop. | -3.16 | 0.002 |
| Winter min SST | 0.993 | 0.479 | 2.70 | 1.21°C | 2.07 | 0.038 |
| Chlorophyll-a | 0.875 | 0.449 | 2.40 | 0.49 mg m^-3^ | 1.95 | 0.051 |
| Summer max SST | 0.764 | 0.477 | 2.15 | 2.20°C | 1.60 | 0.109 |
| Bottom temperature | -0.664 | 0.467 | 0.51 | 2.73°C | -1.42 | 0.155 |
| Park fraction | 0.278 | 0.221 | 1.32 | 0.11 prop | 1.26 | 0.208 |
| Sea ice concentration | -0.880 | 0.750 | 0.41 | 0.045 prop. | -1.17 | 0.241 |
| Salinity | 0.632 | 0.993 | 1.88 | 3.77 PSU | 0.64 | 0.524 |
| Wind speed | 0.192 | 0.381 | 1.21 | 0.47 m s^-1^ | 0.50 | 0.614 |
| Shoreline length | 0.047 | 0.235 | 1.05 | 16 km | 0.20 | 0.843 |
| Precipitation | -0.010 | 0.287 | 0.99 | 14 mm | -0.03 | 0.973 |

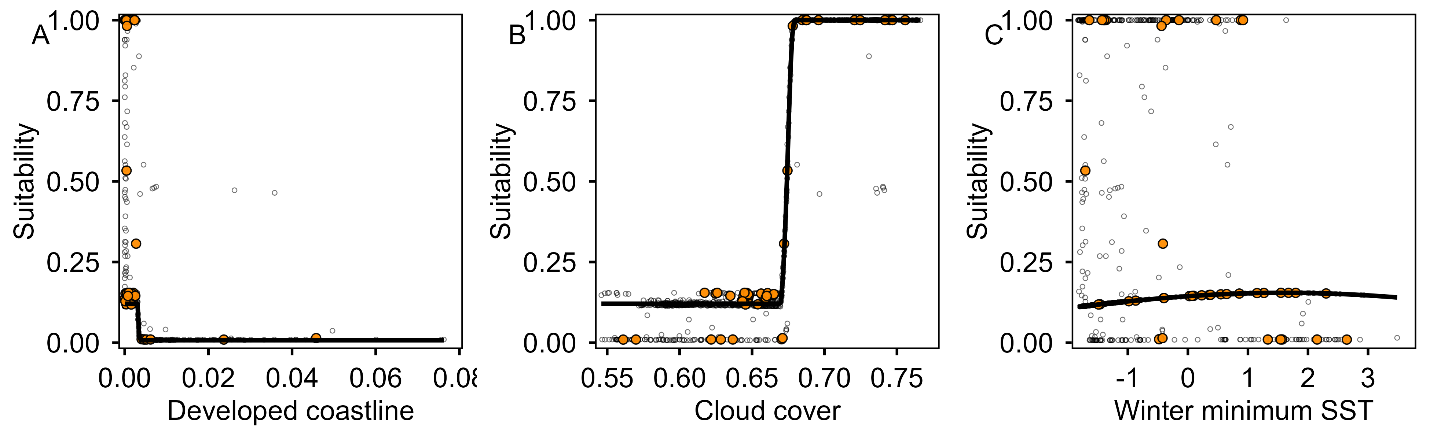

Figure S1. MaxEnt response curves for the three environmental predictors with non-zero coefficients in the final species distribution model. Each panel shows predicted habitat suitability across the observed range of one predictor with all other predictors held at their median value across coastal cells. Coastal cells are overlaid as orange-filled circles for cells with at least one confirmed *Urticina* sp. observation, open circles for cells with no record.

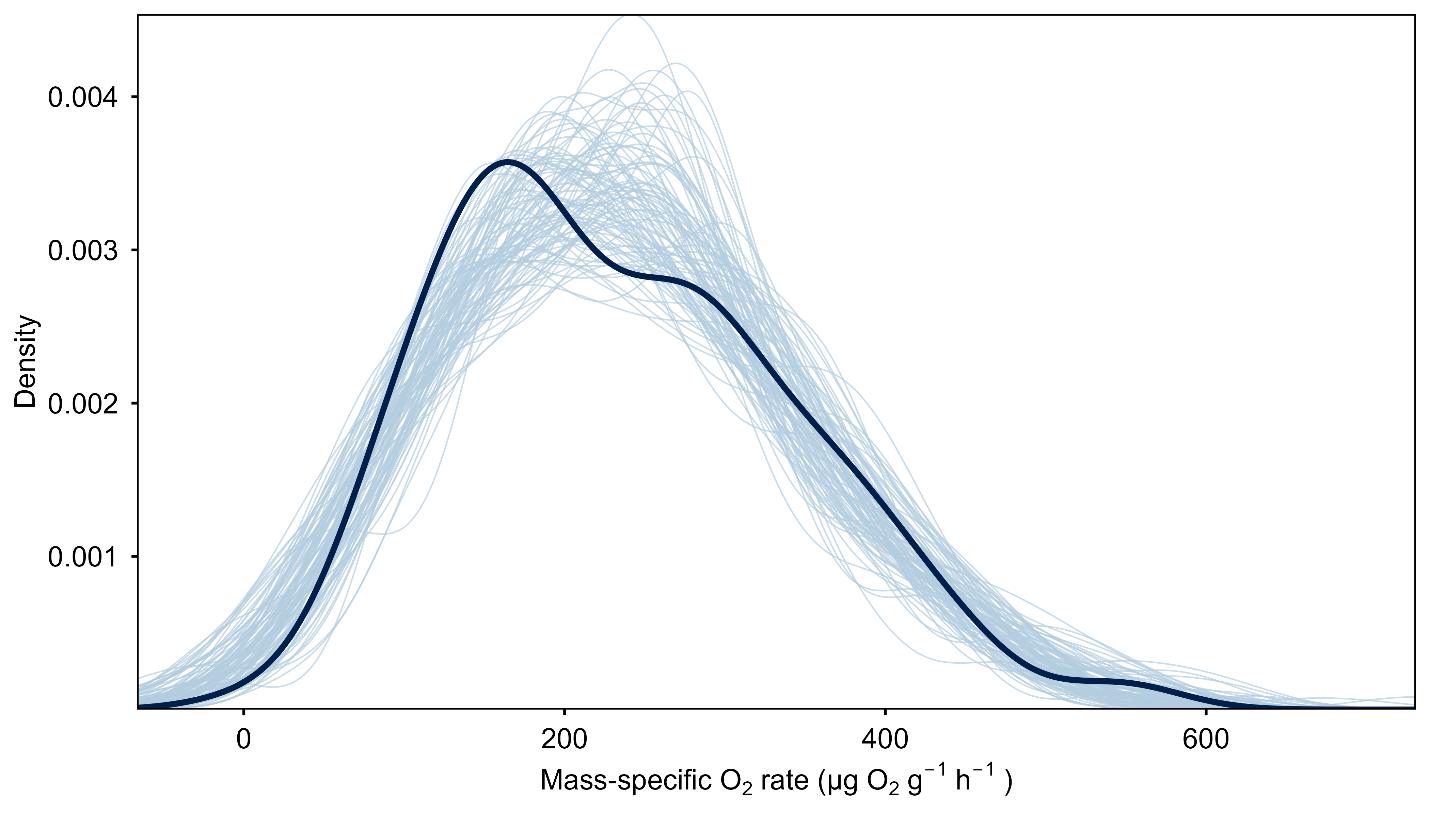

Figure S2. Posterior predictive check for the best-fitting model (Sharpe-Schoolfield Multiplicative). The dark line shows the observed data density; lighter lines show 100 simulated datasets drawn from the posterior predictive distribution. Close agreement indicates that the model adequately captures the distributional properties of the data.

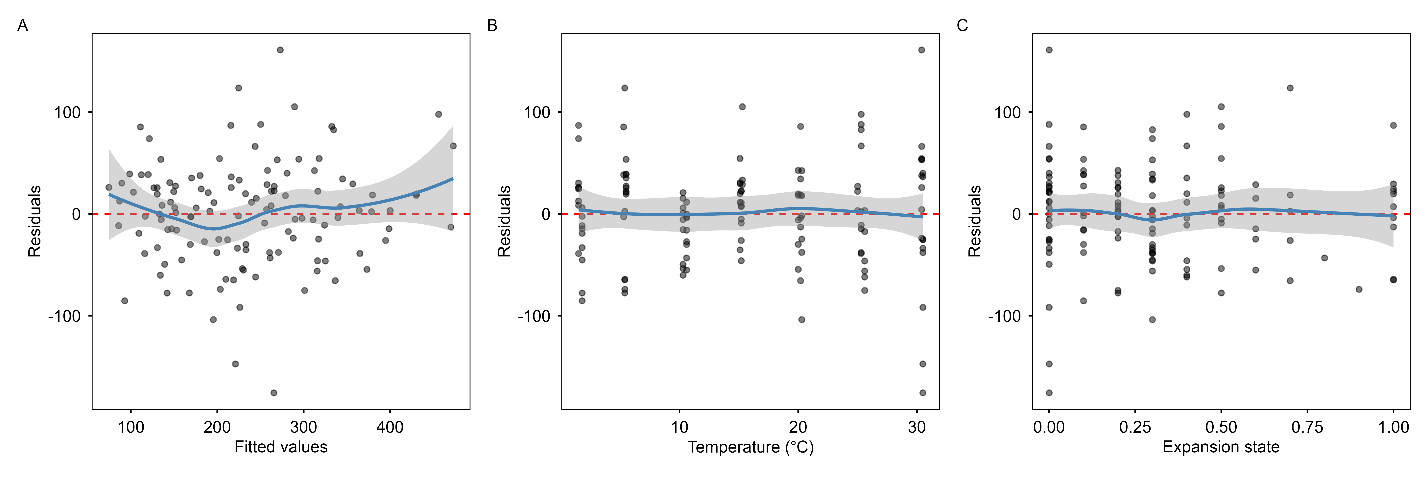

Figure S3. Residual diagnostics for the best-fitting model (Sharpe-Schoolfield Multiplicative). (A) Residuals vs. fitted values; (B) residuals vs. temperature; (C) residuals vs. expansion state. LOESS smooths (blue) indicate no systematic patterns, confirming adequate model specification.

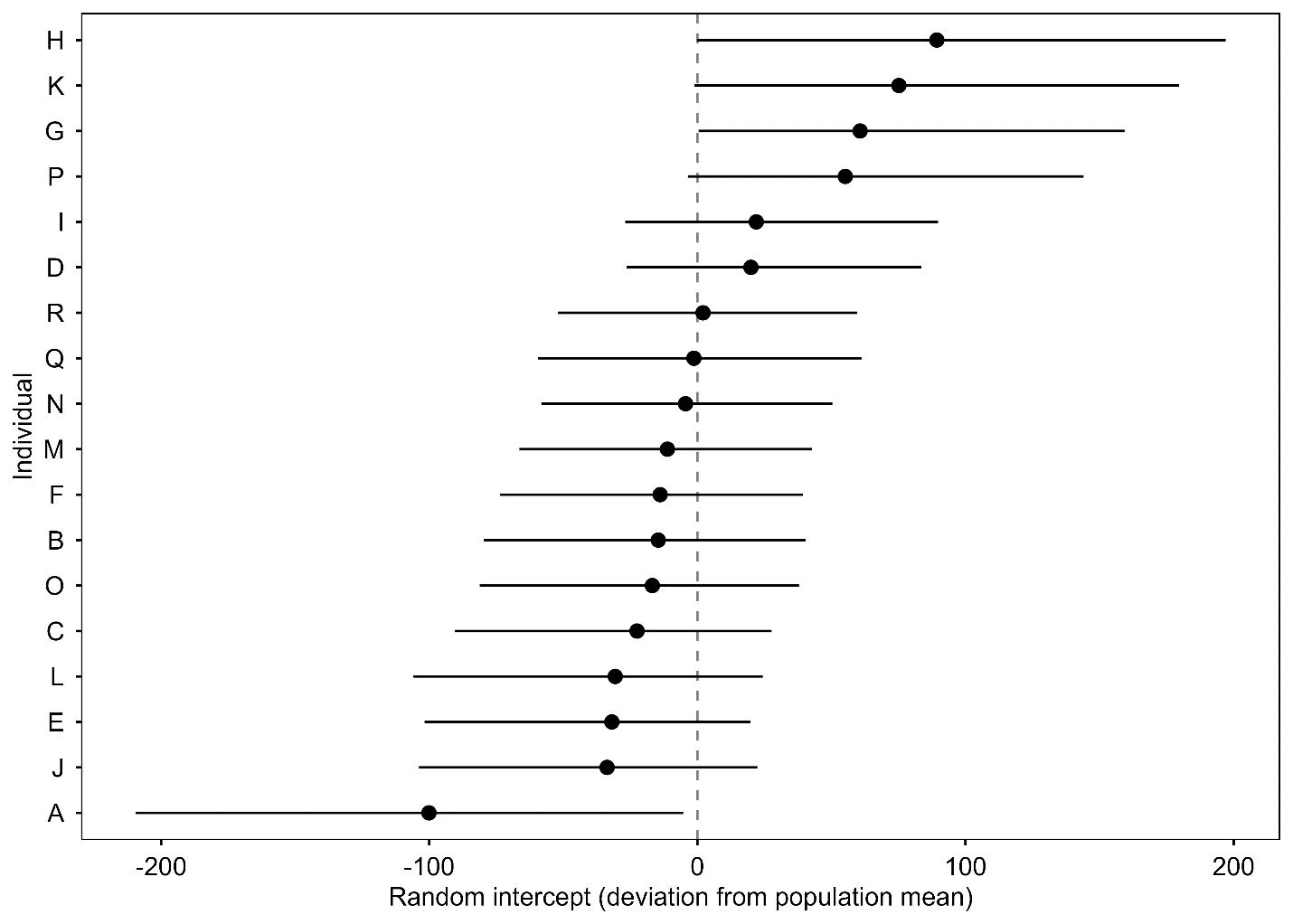

Figure S4. Individual-level random intercept estimates (posterior median and 95% credible interval) for the amplitude parameter of the best-fitting model. Variation across the 18 individuals (relabeled A-R) reflects differences in baseline metabolic rate after accounting for temperature and behavioral state.

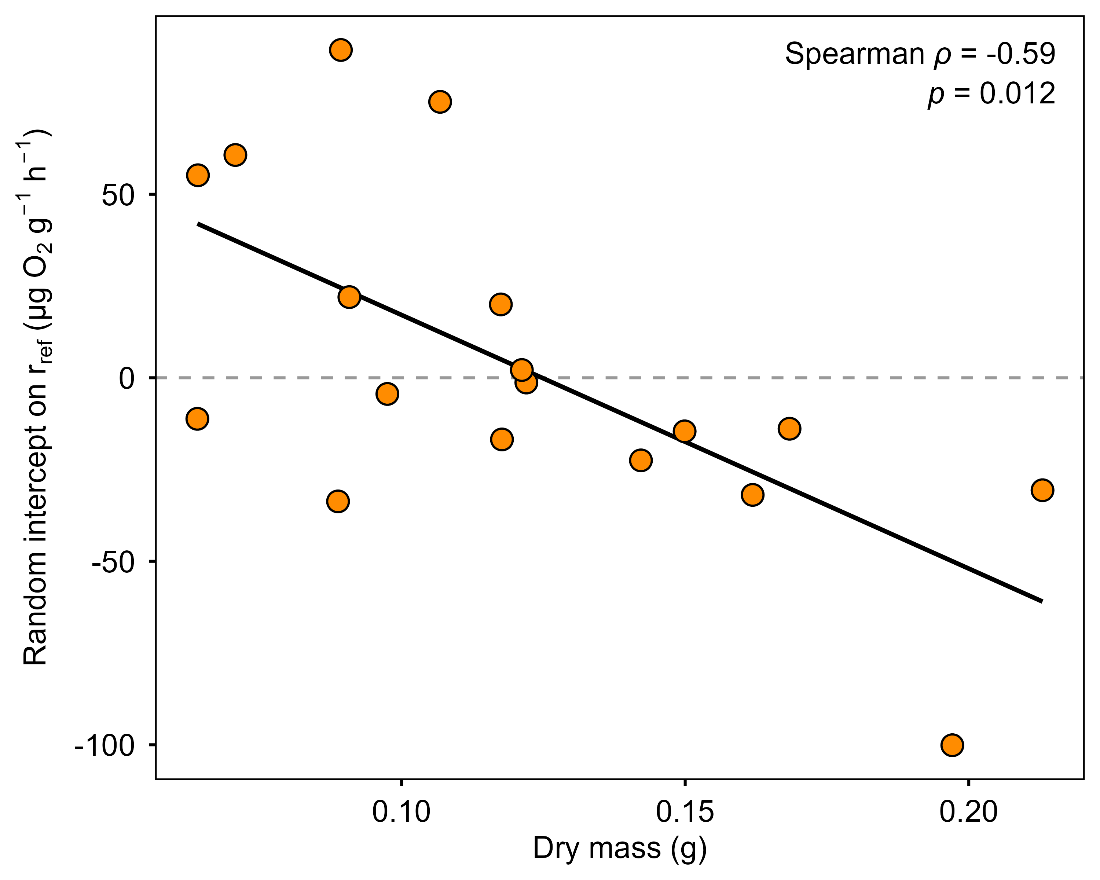

Figure S5. Per-individual random intercept on the reference-temperature rate of the Sharpe-Schoolfield multiplicative model, plotted against dry mass. Each point is one of the 18 anemones; the solid line is a linear least-squares fit.

Table S6. Mass-adjusted thermal performance follow-up. Each behavioral structure (naïve, additive, multiplicative) of the best-performing TPC (Sharpe-Schoolfield) was re-fit with and without the centered logarithm of dry mass as a fixed effect on the model’s amplitude parameter. *β*_mass_ is the posterior median of the dry-mass coefficient with 95% credible intervals in µg O_2_ g^-1^ h^-1^ per log-unit dry mass.

| Structure | Mass adjustment | *T_opt_* (°C) | *CT_max_* (°C) | ΔLOOIC | *β*_mass_ (95% CI) |
| --- | --- | --- | --- | --- | --- |
| Multiplicative | With mass | 26.5 | 45.3 | 0.0 | 16.1 (0.4, 60.8) |
| Multiplicative | Without | 26.5 | 45.9 | 6.0 | — |
| Additive | Without | 25.4 | 45.9 | 8.2 | — |
| Additive | With mass | 25.3 | 46.1 | 8.5 | 18.6 (0.5, 69.1) |
| Naïve | Without | 23.8 | 44.8 | 48.0 | — |
| Naïve | With mass | 23.8 | 45.0 | 48.8 | 15.3 (0.3, 59.3) |
